# Analysis of the CYP1A2 caffeine metabolism gene in the student population at Lake Superior State University

**DOI:** 10.1101/2022.06.14.496190

**Authors:** Rebekka A Pittsley, Stephen H Kolomyjec

**Affiliations:** College of Science and the Environment, Lake Superior State University, Sault Sainte Marie, Michigan, USA

**Author notes:** Corresponding Author:* Rebekka A. Pittsley^*, A^, P.O. Box 291, New Lothrop, MI 48460. Stephen H. Kolomyjec, Lake Superior State University, 650 W. Easterday Ave., Sault Ste. Marie, MI 49783.

**Keywords:** CYP1A2, gene, caffeine, caffeine metabolism

## Abstract

85% of Americans drink caffeinated beverages on a daily basis. Each individual responds differently to caffeine depending on age, gender, diet, and ethnicity. Caffeinated beverages cause insomnia in some people, but not in others due to differences in the rate of caffeine metabolism. This study examines the variation in the caffeine metabolism of Lake Superior State University (LSSU) students. My hypothesis was that LSSU student allele frequencies would match those of the general population: 47.5% fast, 41.0% medium, and 11.5% slow caffeine metabolism. 200 LSSU students were sampled via buccal swabs. DNA was successfully isolated from 164 samples. Participants filled out a demographic questionnaire entailing caffeine intake, ethnicity, and sex. The CYP1A2 gene was amplified via standard PCR prior to genotyping by restriction digest and gel electrophoresis. The APAI restriction enzyme was used to determine the genotype of the rs762551 single nucleotide polymorphism (SNP), while the SACI enzyme was used as a positive digestion control. Overall, results showed a total of 42.7% fast, 44.5% medium, and 12.8% slow metabolizers. Of special note is that 24 of the 164 students sampled were of Native American heritage, an important yet underrepresented group in human genomics. This study provides the first reported look at the CYP1A2 variation within this North American subpopulation with metabolism rates being 50% fast, 33.3% medium, and 16.7% slow. The results confirm my initial hypothesis that the variation of caffeine metabolism gene frequencies for the LSSU student population would be representative of published allele frequencies for the general population.

## Introduction

Caffeine is a white powdery substance with a chemical structure of 1,3,7-trimethylxanthine (Institute of Medicine (U.S.) Committee on Military Nutrition Research (IMCMNR) 2001). Caffeine itself was first isolated in 1819 by Friedlieb Ferdinand Runge and is now recognized as the most used psychoactive drug worldwide (Weinberg *et al.* 2001). In the United States alone, 85% of individuals consume at least one caffeinated drink every day (Mitchell *et al.* 2014). This chemical stimulates the central nervous system, creating a sense of attentiveness once consumed (IMCMNR 2001). Caffeine is easily accessible, inexpensive, and can be beneficial towards brain function (Nikolic *et al.* 2003). Each individual will respond differently to caffeine depending on a variety of factors including age, sex, diet, and Body Mass Index (Nehlig 2018).

As age increases, hepatic liver enzyme function decreases, resulting in an increased sensitivity to caffeine among the elderly (Massey 1998). In contrast, hepatic enzymes are not yet mature in newborns and caffeine will take longer to clear (Nehlig 2018). There is no significant difference among caffeine metabolism between males and females (Nehlig 2018). In general, higher amounts of caffeine consumption take longer to metabolize than smaller amounts. Some people can consume a caffeinated beverage before bed and still fall asleep, while others will be awake for hours. This difference in response is determined by the body’s ability to metabolize caffeine.

The gene that encodes for caffeine metabolism is found on the CYP1A2 gene, which can be found on Chromosome 15, located at the loci 15q24 (Cornelis *et al.* 2011). This gene metabolizes 95% of caffeine consumption, with N-acetyltransferase 2 (NAT2) responsible for roughly the other 5% (NAT2 N-acetyltransferase 2 [Homo sapiens (human)]). The NAT2 enzyme is an acetylator that helps to metabolize drugs and carcinogens. Focusing on CYP1A2, a hepatic cytochrome enzyme, which is responsible for the metabolism of a number of substrates in addition to caffeine that are important in the human body. These substrates include procarcinogens, hormones, drugs, endogenous compounds, as well as enzyme activity in tobacco smokers (Chernyak *et al.* 2011). Procarcinogens like Benzo[a]pyrene, which is a harmless chemical found in many foods like grilled meats and tobacco smoke (Zhou *et al* 2009). Hormones such as estrogen and progesterone, which are sex hormones, are involved in the development of the female reproductive system and in the regulation of the menstrual cycle. Drugs such as Clozapine (schizophrenia medication), Theophylline (asthma medication), as well as Tylenol (pain medication). Endogenous compounds are substances that originate from inside the human body. Examples of these include steroids and arachidonic acid (polyunsaturated fatty acid present in phospholipids). Major substrate classes including irritable bowel syndrome (Alosetron), Estrogen (Estradiol), and Anti-Parkinson: dopamine agonist (Ropinirole) to name a few (Fankhauser 2013). Minor substrate classes include, but are not limited to, medications such as Acetaminophen, Melatonin, Warfarin, and Progesterone (Fankhauser 2013).

In the late 1990’s, various researchers began analyzing the base pair substitution from A to C in the CYP1A2 gene to determine its importance. A German study by Sachase *et al.* (1999) concluded that the base pair changes led to polymorphism variations, with the AA genotype leading to increased gene activity. In 2007, a Canadian study pinpointed the differing rates of caffeine metabolism related to variation in the gene (El-Sohemy *et al.* 2007). It was noted that a base pair substitution of A to C at a key point in the CYP1A2 gene changes the metabolism rate. The homozygous allele (AA) metabolizes caffeine at a fast rate, the heterozygous allele (CA) metabolizes caffeine at a medium rate, and the homozygous allele (CC) metabolizes caffeine at a slow rate (Zephyr and Walsh 2015).

A 2012 meta-analysis of 8,345 Caucasians showed the prevalence of the CYP1A2 gene variations. This study’s control population found that 50.3% Caucasians were fast metabolizers (AA), 41.5% were medium metabolizers (CA), and 8.2% were slow metabolizers (CC). The 2,423 people of Asian descent sampled showed 40.8% AA, 44.8% CA, and 14.4% CC (Wang *et al.*, 2012). A Japanese study (Shimada *et al.* 2009) of 403 Japanese nationals noted 40.4% AA, 46.2% CA, and 13.4% CC, while 389 Brazilians exhibited an allele frequency of 45.0% AA, 41.1% CA, and 13.9% CC.

The single nucleotide polymorphism (SNP) that determines the differing levels of enzyme function can be determined in multiple ways such as through high throughput SNP assay chips, medium throughput qPCR probes, or lower throughout restriction fragment length polymorphism (RFLP). The SNP in question occurs within a known restriction site (Zephyr and Walsh 2015), the results of a simple restriction digest can be used to determine which SNP is present. This particular SNP (rs762551) is in an intron of DNA that can increase transcription rates and increase mRNA translation efficiency (Shaul 2017). The CYP1A2 gene consists of two alleles. Whether there is a polymorphism of A or C base pair at this snip will determine the metabolism rate at which caffeine is processed.

Restriction fragment length polymorphism (RFLP) is recognized by the restriction enzyme and cuts the DNA at specific sites. RFLP markers isolate as codominant alleles, allowing for the comparison of genetic structure parameters using genetic variability (Yan *et al.* 1999). Depending on the allele present, in this case the A or C on the CYP1A2 gene, the enzyme will or will not cut at that site, resulting in three various fragment lengths: 249 bp, 494 bp, and 743 bp (Figure 2).

Chi-square tests were used to compare and contrast the results from the expected results. Expected results of allele frequencies were used from the 1000 Genomes Project Phase 3 (Ensembl 2021). Subpopulation frequencies used included African, Eastern Asian, European, Finnish, and British frequencies.

The objective of this study was to examine the variation in the caffeine metabolism of Lake Superior State University (LSSU) students. We hypothesized that the LSSU student allele frequencies would match those of the European population, since a majority of the students are of European descent. Those expected allele frequencies being 47.5% fast (AA), 41.0% medium (AC), and 11.5% slow (CC) caffeine metabolizers (Ensembl 2021).

## Methods

Lloyd, 2003 analyzed the C677T mutation in the MTHFR gene and employed similar methods that were used in the analysis of the CYP1A2 gene. The study population, DNA extraction, PCR amplification, restriction digest, and fragment analysis sections of Lloyd’s paper were used and adapted for the general methods and procedure of this study.

### Study Population

To determine the variation of potential caffeine sensitivity within a local population, two-hundred DNA samples were collected from students at LSSU in Sault Ste. Marie, Michigan. Samples were numbered one through two-hundred and obtained by cheek swabs. All participation was on a voluntary basis with IRB approval #05292020. The samples were collected by visiting classrooms of participating professors and explaining my project. Each subject received and signed a copy of the consent form approved through the Institutional Review Board for the Protection of Human Subjects at LSSU. Students also filled out an anonymous demographic questionnaire to accompany the study detailing questions regarding age, gender, ethnicity, and caffeine consumption. Each student provided only one sample for this study. Students were all at least 18 years of age and active full-time or part-time LSSU students.

Each participant was asked to gently scrape the inside of their own cheek using a sterile mouth swab. The swab was then given to the primary investigator and placed in a 1.5 ml microcentrifuge tube containing 400 μl of Cell Lysis Buffer from the Monarch Genomic DNA Purification Kit (NEB). Samples were stored in Cell Lysis Buffer at 4°C until processed.

### DNA Extraction

Genomic DNA was isolated using the Monarch Genomic DNA Purification Kit (NEB) Sample Lysis: Animal Tissue protocol. The modifications to the general procedure are as follows: each sample contained 400 ul of Cell Lysis Buffer and mouth swab; 3 μl of RNase A was added to the lysate of samples, excluding #80, 86-103 due to lack of supplies.

### PCR Amplification

Polymerase Chain Reaction (PCR) amplification was performed using published primers for the CYP1A2 gene (Table 2) and both the VWR Life Science Hot start PCR-to-Gel Taq Master mix, 2X and the Thermo Science DreamTaq Green DNA Polymerase (5 U/μL). The total volume of each reaction was 25 μl and consisted of 12 μl master mix, 2 μl primer, 8 μl deionized water, and 5 μl genomic DNA. This reaction was multiplied by three, to allow for multiple restriction digests, for a total reaction volume of 81 μl.

**Table 1.** Metabolism rates according to population and subpopulation. All values less than the critical Chi Square: 3.841. Not significantly different from published allele frequencies.

| **Population** | **Sample Size** | **Fast** | **Actual**<br>**Medium** | **Slow** | **Experimental**<br>**$\chi^2$ Value** |
| --- | --- | --- | --- | --- | --- |
| European | 139 | 43.8% | 43.2% | 12.9% | 0.3914 |
| African American | 8 | 25% | 50% | 25% | 0.2447 |
| Asian | 7 | 12.3% | 57.1% | 28.6% | 3.786 |
| Native American | 24 | 50% | 33.3% | 16.7% | 0.923 |
| Finnish | 11 | 54.5% | 33.3% | 18.2% | 0.9642 |
| British | 28 | 53.6% | 39.3% | 7.1% | 0.1651 |

**Table 2.**
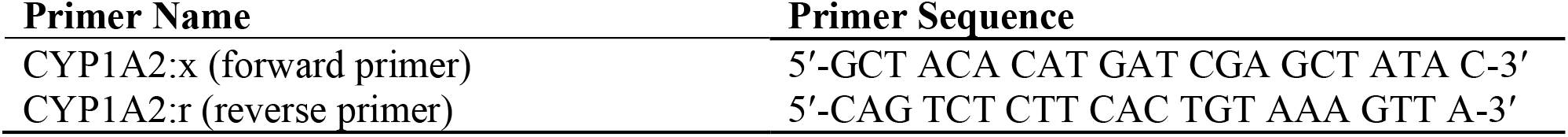
Primers used in the PCR amplification of the CYP1A2 gene (Eshkoor et. al 2013).

**Table 3.** Chi Square composing project allele frequencies from by ethnicity (Ensemble, 2021). Critical Chi Square: all populations failed to reject the null hypothesis at alpha = 0.05.

| European | Allele | Actual | Expected |  |  |
| --- | --- | --- | --- | --- | --- |
|  | AA | 61 | 66 | X2= | 0.3914 |
|  | AC | 60 | 57 |  |  |
|  | CC | 18 | 16 |  |  |
|  | Total | 139 | 139 |  |  |
| African American | Allele | Actual | Expected |  |  |
|  | AA | 2 | 2.576 | X2= | 0.2774 |
|  | AC | 4 | 3.84 |  |  |
|  | CC | 2 | 1.584 |  |  |
|  | Total | 8 | 8 |  |  |
| Asian | Allele | Actual | Expected |  |  |
|  | AA | 1 | 3.178 | X2= | 3.788 |
|  | AC | 4 | 3.059 |  |  |
|  | CC | 2 | 0.763 |  |  |
|  | Total | 7 | 7 |  |  |
| Native American* | Allele | Actual | Expected |  |  |
|  | AA | 12 | 11.4 | X2= | 0.9327 |
|  | AC | 8 | 9.84 |  |  |
|  | CC | 4 | 2.76 |  |  |
|  | Total | 24 | 24 |  |  |
| Finland | Allele | Actual | Expected |  |  |
|  | AA | 6 | 5.445 | X2= | 0.9642 |
|  | AC | 3 | 4.334 |  |  |
|  | CC | 2 | 1.221 |  |  |
|  | Total | 11 | 11 |  |  |
| England | Allele | Actual | Expected |  |  |
|  | AA | 15 | 15.372 | X2= | 0.1651 |
|  | AC | 11 | 10.164 |  |  |
|  | CC | 2 | 2.464 |  |  |
|  | Total | 28 | 28 |  |  |

\*Compared to European allele frequencies

**Table 4.** European Breakdown Statistics of caffeine metabolism variance across geographic region.

| **Geographic**<br>**Region** | **Northern**<br>**Europe** | **Eastern**<br>**Europe** | **Central**<br>**Europe** | **Western**<br>**Europe** | **Southern**<br>**Europe** |
| --- | --- | --- | --- | --- | --- |
| AA | 53% | 66% | 8% | 24% | 23% |
| AC | 35% | 16% | 63% | 52% | 38% |
| CC | 10% | 16% | 28% | 22% | 38% |

Amplification was completed using a Veriti Thermal Cycler (Applied Biosystems International) using the parameters described by Zephyr and Walsh (2015). The PCR reaction occurred as follows: 1 cycle of denaturation at 95°C for 5 minutes, followed by 35 cycles at 95°C for 30 seconds, 57.5°C for 30 seconds, and 68°C for 1 minute. The final stage had 1 cycle occur at 68°C for 5 minutes. Post-PCR product was then refrigerated and stored at 4°C (Zephyr and Walsh 2015).

### Restriction Digest

The restriction enzymes used to digest the Post-PCR product were NEB CutSmart APAI and SACI. The APAI enzyme was used to detect the SNP and did not digest alleles containing A, which resulted in 743 bp, but APAI did digest alleles containing C (494 bp and 249 bp). The SACI enzyme served as the positive control restriction digest with base pairs appearing in fragment analysis at 249 and 494.

PCR products underwent restriction digest with a total volume of 50 μl each. Each restriction digest consisted of 24 μl H_2_O, 20 μl of PCR DNA product, 5 μl CutSmart Buffer, and 1 μl of each restriction enzyme: APAI or SACI. The restriction digests were set up in 200 μl PCR tubes. The tubes were flicked gently to mix and centrifuged briefly. The SACI enzyme was incubated at 37°C for 1 hour and the APAI enzyme was incubated at 25°C for 1 hour. 10 μl of Purple Gel Loading dye was added to each 50 μl SACI reaction to stop reaction.

Results were visualized and genotyped via agarose gel electrophoresis using 1.5% gels pre-stained with GelRed (Biotium). 10 μl of DNA ladder, 15 μl of PCR product, 15-20 μl of the APAI samples, and 15 μl of the negative control SACI were added to the wells.

### Data Collection & Analysis

Data was collected and each sample was classified as either homozygous uncut (AA), heterozygous (AC), or homozygous cut (CC), based on the length of specific base pair fragments. Homozygous uncut (AA) had two A alleles with two base pair lengths at 743. Heterozygous (AC) had one A allele and one C alle with base pair lengths at 743 (A), 494 bp, and 249 bp respectively (C). Homozygous cut (CC) had 2 C alleles with two sets of base pair lengths at 494 and 249 (C). A Chi-squared test with 5% percent deviation (χ^2^ value: 3.841) was used to determine the accuracy of the collected data in comparison to published frequencies on Ensemble.

## Results

200 samples were collected from students. Of that, DNA was successfully extracted from 164 of the samples. The 36 samples that did not work were re-run from PCR with no results a second time. Of the 164 samples, ethnicities were broken down into the major population types: European, African American, Asian, and Native American, with European sub-populations of comparable published allele frequencies of Finnish and British. For the European population there was 139 students total with metabolisms of 43.8% fast, 43.2% medium, and 12.9% slow. The experimental χ^2^ value for Europeans was 0.3914. For the African American population there was 8 students total with metabolisms of 25% fast, 50% medium, and 25% slow. The experimental χ^2^ value for African Americans was 0.2447. For the Asian population there was 7 students with metabolisms of 12.3% fast, 57.1% medium, and 28.6% slow. The experimental χ^2^ value for Asians was 3.786. For the Native American population there was 24 students total with metabolisms of 50% fast, 33.3% medium, and 16.7% slow. For the European sub-population of Finnish, there were 11 students with metabolisms of 54.5% fast, 33.3% medium, and 18.2% slow. The experimental χ^2^ value for Finnish was 0.9642. For the European sub-population of British, there were 28 students with metabolisms of 53.6% fast, 39.3% medium, and 7.1% slow. The experimental χ^2^ value for the British population was 0.1651. All χ^2^ values were under 3.841, showing that the data was not significantly different from the published allele frequencies.

Looking closer at the breakdown of the European regions, the overall European caffeine metabolism was 43.8% fast, 43.2% medium, and 12.9% slow. The caffeine metabolism of each region varied (Figure 1, Table 4). The Northern European demographic includes Finland, Denmark, Sweden, and Norway. Of the 21 total individuals, 53% were fast metabolizers, 35% were medium metabolizers, and 10% were slow metabolizers. The Eastern European demographic includes Ukraine, Russia, and Armenia. Of the 5 total individuals, 66% were fast metabolizers, 16% were medium metabolizers, and 16% were slow metabolizers. The Central European demographic includes Germany, Switzerland, Poland, Hungary, Austria, and Slovakia. Of the 107 total individuals, 11% were fast metabolizers, 68% were medium metabolizers, and 21% were slow metabolizers. The Western European demographic includes Belgium, France, Germany, Switzerland, Netherlands, Czech Republic, Ireland, England, and Scotland. Of the 171 total individuals, 24% were fast metabolizers, 52% were medium metabolizers, and 22% were slow metabolizers. The Southern European demographic includes Spain, Turkey, Italy. Of the 18 total individuals, 23% were fast metabolizers, 38% were medium metabolizers, and 38% were slow metabolizers. In summary, Northern and Eastern Europe had the highest number of fast caffeine metabolizers, Central and Western Europe had mostly medium metabolizers, and Southern Europe had an equal distribution of medium and slow metabolizers.

**Figure 1:**
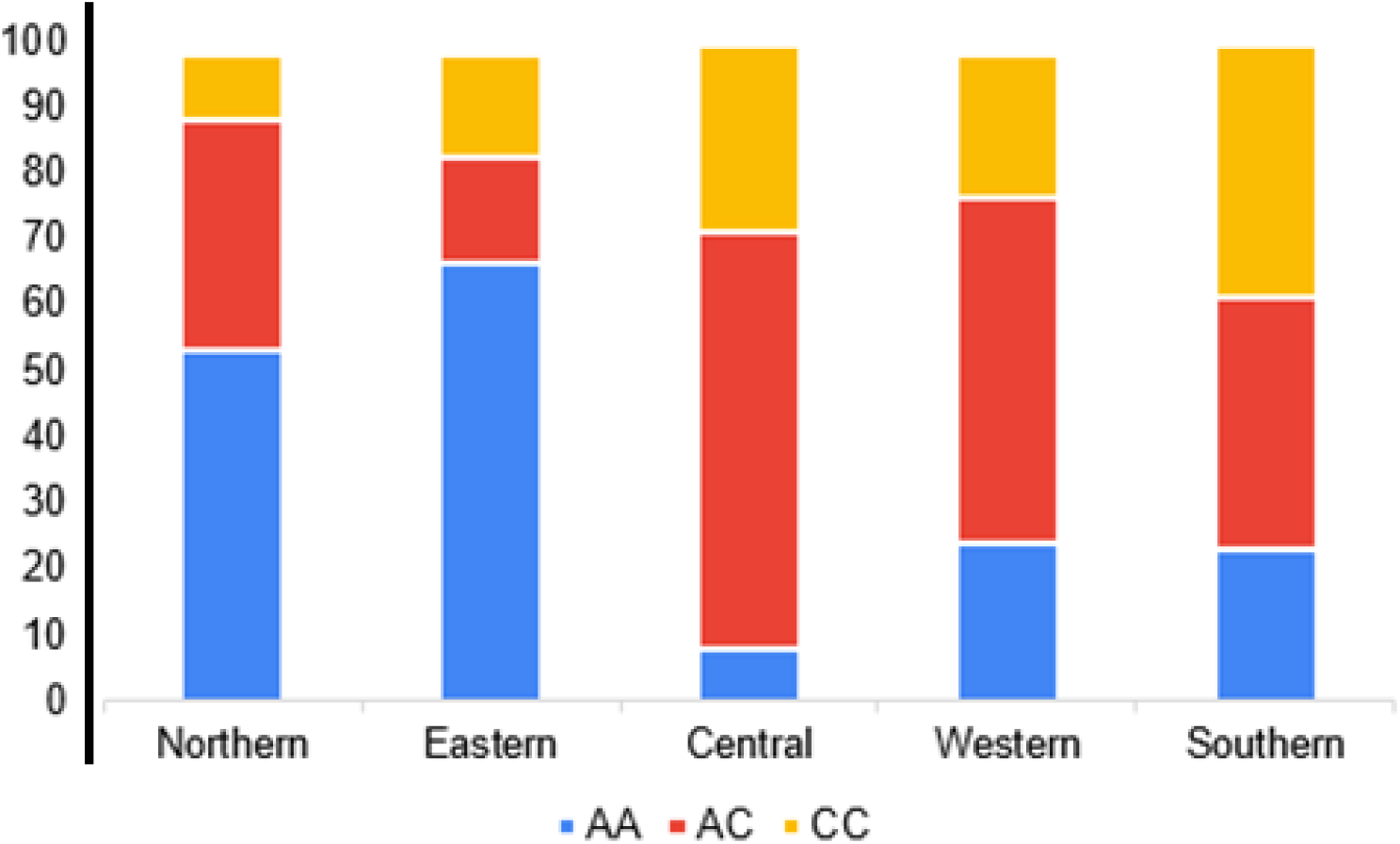
Breakdown of Europe into geographic regions. Caffeine metabolism varies across European geographic distribution.

**Figure 2.**
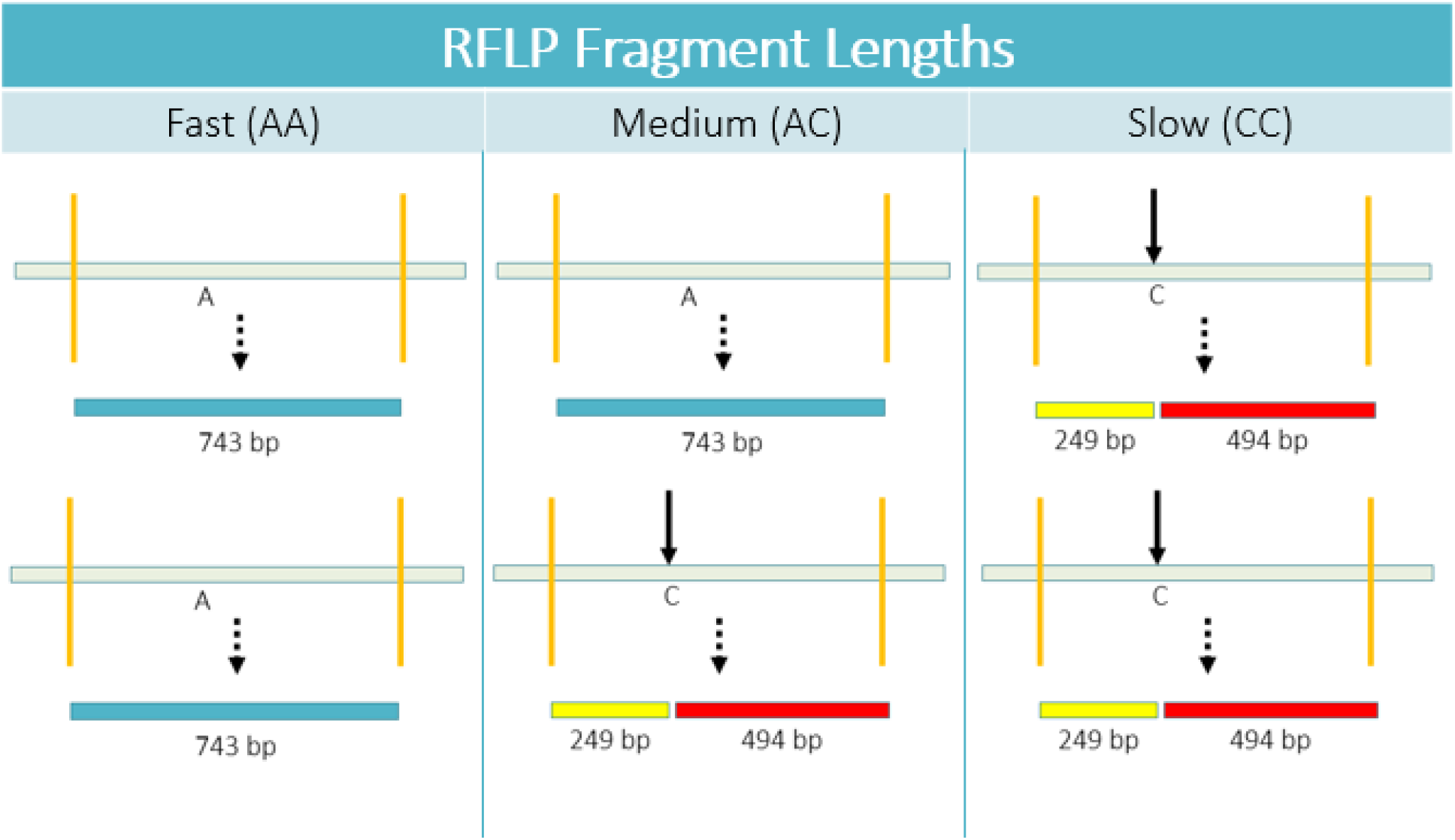
Restriction Fragment Length Polymorphisms: Fast, Medium, and Slow Fragment Lengths.

There were 24 individuals that were of Native American heritage, comprising 14.6% of the population. This population is understudied and I was not able to find any comparative previously published data. In order to run the χ^2^ test, the expected European χ^2^ value was used to compare the observed Native American data. The experimental χ^2^ value was 0.923. The African American population had an interesting caffeine metabolism population. It showed 50% fast metabolizers, 33.3% medium, and 16.7% slow metabolizers. It had a larger population of fast metabolizers, a smaller population of medium metabolizers, and a larger population of slow metabolizers than the European population.

For sex, there was no significant difference and the data matched between the two groups – 42.5% of females and 43.4% of males were fast metabolizers; 45.2% of females and 43.3% of males were medium metabolizers, and 12.3% of females and 13.3% of males were slow metabolizers.

For age, a significant amount of the population were of the Gen Z demographic (born 1997 to 2015), with 152 of the 164 samples (92.7%) being in the age range of 18 to 24. 43.4% were fast metabolizers, 44.1% were medium metabolizers, and 13.2% were slow metabolizers.

## Discussion

The Chi-Square Test results confirmed the hypothesis: LSSU students sampled were representative of the European population, as their allele frequencies did not significantly vary from the published frequencies. Predicted allele frequencies were 47.5% fast, 41.0% medium, and 11.5% slow (Ensembl 2021). The actual allele frequencies of LSSU students were 42.7% fast, 44.5% medium, and 12.8% slow. European, African American, Asian, Native American, Finnish, and British population experimental Chi-Square values were less than the critical Chi Square value of 3.841. Thus, the major population and the subpopulations at LSSU are not significantly different from the general population. Age and sex also had no significant differences from the European allele frequencies, as the CYP1A2 gene is not sex-linked.

The data gathered from the 7 Asian students showed a majority of medium metabolizers (57.1%), second-most slow metabolizers (28.6%), and the fewest fast metabolizers (14.3%). The experimental χ^2^ value was 3.786. This value is less than the critical χ^2^ value of 3.841, but not by much. This gathered data was compared to the published Asian allele frequencies and were found to be 32% fast metabolizers, 31% medium metabolizers, and 8% slow metabolizers. One meta-analysis of 2,423 people of Asian descent showed caffeine metabolism levels of 40.8% AA, 44.8% AC, and 14.4% CC (Wang *et al.* 2012). The difference between the populations can be contributed to sample size variations. For the LSSU population, there was sample size of 2 fast metabolizers, 7 medium metabolizers, and 4 slow metabolizers. Based on the Ensemble sample size, there was 229 fast metabolizers, 220 medium metabolizers, and 55 slow metabolizers. The meta-analysis study (Wang *et al.* 2012) had 989 fast metabolizers, 1086 medium metabolizers, and 348 slow metabolizers.

The European distribution of caffeine metabolism throughout the different regions showed remarkable differences between Sault Ste. Marie populations, some of this could be due in part to sample size. This showed that the caffeine metabolism was dependent on geographic location, as Northern and Eastern Europe had the most fast caffeine metabolizers, Central and Western Europe had the most medium metabolizers, and Southern Europe had an equal distribution of medium and slow metabolizers.

Of special note is that 24 of the 164 students sampled were of Native American heritage, an important yet underrepresented group in human genomics. This study provides the first reported look at the CYP1A2 variation within this North American subpopulation with metabolism rates being 50% fast, 33.3% medium, and 16.7% slow. In order to run the χ^2^ test, the expected European χ^2^ value was used to compare the observed Native American data. The experimental χ^2^ value was 0.923. It was noted that 19 of the 24 Native American population also had an overlapping European heritage. Thus, the Native American results are not significantly different from the European results.

Knowing your caffeine metabolism could improve your health. Consuming too much caffeine can cause negative side effects such as: increased blood pressure, insomnia, heart palpitations, dehydration, headaches, nervousness, irritability, and muscle tremors (Mayo Clinic 2020). Slow metabolizers should avoid excessive caffeine consumption as they are at risk for heart attacks (El-Sohemy *et al.* 2007). There is the possibility of interactions with medications and supplements. Such medications interactions include Quinolones antibiotics, which decrease caffeine metabolism, and Bronchodilators, which are also stimulants (Mayo Clinic 2020). Other medications include Tylenol, which the CYP1A2 is a minor substrate metabolizer. In turn, an individual who takes longer to metabolize these substrates will experience the drug longer and is at risk for various side effects (Fankhauser 2013).

The focus of this study was college students and looked at their caffeine metabolism, their caffeine consumption, and how their ethnicity plays a role in their metabolism. College students are known to consume large amounts of caffeine – whether it be drinking an energy drink to pull an all-nighter to finish a paper or a coffee in the morning to get going. Slow, medium, and fast metabolizers will have to consume different amounts of caffeine to ‘wake-up’ or get a caffeine buzz. For example, a fast metabolizer might have to get a coffee with two shots of espresso whereas a slow metabolizer might have consumed the same drink and not be able to sleep until 2AM. I would warn the slow metabolizers to be careful when consuming excess amounts of caffeine because they are more at risk for heart attacks or to experience previously mentioned side effects like jitteriness or heart palpitations (Mayo Clinic 2020). One should also take into consideration that the hepatic liver enzyme function decreases as age increases, resulting in an increased sensitivity to caffeine (Massey 1998). Thus, someone who was able to down a 5-Hour Energy in college right before bed might not be able to do the same when they are older.

## Conclusion

The allele frequency of caffeine metabolizers in Lake Superior State University (LSSU) students was determined. Gathered allele frequencies did not significantly differ from published allele frequencies. Predicted allele frequencies were 47.5% fast, 41.0% medium, and 11.5% slow. The actual allele frequencies of LSSU students were 42.7% fast, 44.5% medium, and 12.8% slow. The Chi-Square Test results confirmed the hypothesis: LSSU students sampled were representative of the European population, as their allele frequencies did not significantly vary from the published frequencies.

## Acknowledgements

We would like to thank the following: Drs. Britton Ranson-Olson and Martha Hutchens for their assistance with protocol development; Lake Superior State University for lab space and infrastructure; the LSSU IRB Committee for providing oversight and allowing the research to occur (IRB approval #05292020); the LSSU Undergraduate Research Committee for funding the project; and the students that helped with their time and support - Haven Borghi, Hannah Sawicki, and Jacob Pittsley.

## Data Availability

Raw genotyping data is available in the Supplemental File S2.

## Supplemental Data A

|  | AA | AC | CC | XX | Total Samples |
| --- | --- | --- | --- | --- | --- |
| Total Sample Gathered | 70 | 73 | 21 | 36 | 200 |
| Successful DNA Extraction of Samples | 70 | 73 | 21 |  | 164 |
| Overall Percentage | 42.68 | 44.51 | 12.80 |  |  |

| Ethnicity | AA | AC | CC | Total | Percentage |  |  |
| --- | --- | --- | --- | --- | --- | --- | --- |
|  |  |  |  |  | AA | AC | CC |
| European | 61 | 60 | 18 | 139 | 43.88 | 43.17 | 12.95 |
| African American | 2 | 4 | 2 | 8 | 25.00 | 50.00 | 25.00 |
| Asian | 1 | 4 | 2 | 7 | 14.29 | 57.14 | 28.57 |
| Native American | 12 | 8 | 4 | 24 | 50.00 | 33.33 | 16.67 |
| African | 1 | 1 | 0 | 2 | 50.00 | 50.00 | 0.00 |
| Mexicans | 1 | 1 | 0 | 2 | 50.00 | 50.00 | 0.00 |
| **Sex** |  |  |  |  | **AA** | **AC** | **CC** |
| Female | 31 | 33 | 9 | 73 | 42.47 | 45.21 | 12.33 |
| Male | 39 | 39 | 12 | 90 | 43.33 | 43.33 | 13.33 |
| Unidentified | 0 | 1 | 0 | 1 | 0.00 | 100.00 | 0.00 |
| **Age** |  |  |  |  | **AA** | **AC** | **CC** |
| Gen Z (1997-2015) | 66 | 67 | 20 | 152 | 43.42 | 44.08 | 13.16 |
| Millennials (1981-1996) | 3 | 6 | 1 | 10 | 30.00 | 60.00 | 10.00 |
| Generation X (1965-1980) | 1 | 0 | 0 | 1 | 100.00 | 0.00 | 0.00 |

| Breakdown by<br>European Nationality | AA | AC | CC | Total | Percentage |  |  |
| --- | --- | --- | --- | --- | --- | --- | --- |
|  |  |  |  |  | AA | AC | CC |
| Denmark | 3 | 0 | 0 | 3 | 100.00 | 0.00 | 0.00 |
| England | 15 | 11 | 2 | 28 | 53.57 | 39.29 | 7.14 |
| Finland | 6 | 3 | 2 | 11 | 54.55 | 27.27 | 18.18 |
| France | 3 | 3 | 6 | 12 | 25.00 | 25.00 | 50.00 |
| Germany | 22 | 39 | 8 | 69 | 31.88 | 56.52 | 11.59 |
| Ireland | 19 | 14 | 5 | 38 | 50.00 | 36.84 | 13.16 |
| Italy | 5 | 7 | 2 | 14 | 35.71 | 50.00 | 14.29 |
| Netherlands | 1 | 6 | 1 | 8 | 12.50 | 75.00 | 12.50 |
| Norway | 1 | 2 | 0 | 3 | 33.33 | 66.67 | 0.00 |
| Poland | 11 | 17 | 4 | 32 | 34.38 | 53.13 | 12.50 |
| Scandinavian | 11 | 7 | 3 | 21 | 52.38 | 33.33 | 14.29 |
| Scotland | 6 | 5 | 1 | 12 | 50.00 | 41.67 | 8.33 |
| Sweden | 1 | 2 | 1 | 4 | 25.00 | 50.00 | 25.00 |
| Average |  |  |  |  | 42.95 | 42.67 | 14.38 |

| **Northern Europe** | **AA** | **AC** | **CC** | **Total** | **AA** | **AC** | **CC** |
| --- | --- | --- | --- | --- | --- | --- | --- |
| Denmark | 3 | 0 | 0 | 3 | 100.00 | 0.00 | 0.00 |
| Finland | 6 | 3 | 2 | 11 | 54.55 | 27.27 | 18.18 |
| Norway | 1 | 2 | 0 | 3 | 33.33 | 66.67 | 0.00 |
| Sweden | 1 | 2 | 1 | 4 | 25.00 | 50.00 | 25.00 |
| **Eastern Europe** | **AA** | **AC** | **CC** | **Total** | **AA** | **AC** | **CC** |
| Armenia | 1 | 0 | 0 | 1 | 100.00 | 0.00 | 0.00 |
| Russia | 1 | 0 | 1 | 2 | 50.00 | 0.00 | 50.00 |
| Ukraine | 1 | 1 | 0 | 2 | 50.00 | 50.00 | 0.00 |
| **Central Europe** | **AA** | **AC** | **CC** | **Total** | **AA** | **AC** | **CC** |
| Austria | 0 | 1 | 0 | 1 | 0.00 | 100.00 | 0.00 |
| Germany | 22 | 39 | 8 | 69 | 31.88 | 56.52 | 11.59 |
| Hungary | 0 | 1 | 1 | 2 | 0.00 | 50.00 | 50.00 |
| Poland | 11 | 17 | 4 | 32 | 34.38 | 53.13 | 12.50 |
| Slovakia | 0 | 1 | 1 | 2 | 0.00 | 50.00 | 50.00 |
| Switzerland | 0 | 1 | 0 | 1 | 0.00 | 100.00 | 0.00 |
| **Western Europe** | **AA** | **AC** | **CC** | **Total** | **AA** | **AC** | **CC** |
| Belgium | 0 | 2 | 0 | 2 | 0.00 | 100.00 | 0.00 |
| Czech Republic | 0 | 0 | 1 | 1 | 0.00 | 0.00 | 100.00 |
| England | 15 | 11 | 2 | 28 | 53.57 | 39.29 | 7.14 |
| France | 3 | 3 | 6 | 12 | 25.00 | 25.00 | 50.00 |
| Germany | 22 | 39 | 8 | 69 | 31.88 | 56.52 | 11.59 |
| Ireland | 19 | 14 | 5 | 38 | 50.00 | 36.84 | 13.16 |
| Netherlands | 1 | 6 | 1 | 8 | 12.50 | 75.00 | 12.50 |
| Scotland | 6 | 5 | 1 | 12 | 50.00 | 41.67 | 8.33 |
| Switzerland | 0 | 1 | 0 | 1 | 0.00 | 100.00 | 0.00 |
| **Southern Europe** | **AA** | **AC** | **CC** | **Total** | **AA** | **AC** | **CC** |
| Italy | 5 | 7 | 2 | 14 | 35.71 | 50.00 | 14.29 |
| Spain | 1 | 2 | 0 | 3 | 33.33 | 66.67 | 0.00 |
| Turkey | 0 | 0 | 1 | 1 | 0.00 | 0.00 | 100.00 |

